# A novel fishing cat reference genome for the evaluation of potential germline risk variants

**DOI:** 10.1101/2022.11.17.516921

**Authors:** Rachel A. Carroll, Edward S. Rice, William J. Murphy, Leslie A. Lyons, Lyndon Coghill, William F. Swanson, Karen A. Terio, Tyler Boyd, Wesley C. Warren

## Abstract

The fishing cat, *Prionailurus viverrinus*, displays a fish hunting behavior uncommon among most other cats. Estimated population declines in the wild increase the significance of its existing zoo populations, particularly with a recent high prevalence of transitional cell carcinoma (TCC), a form of bladder cancer. We hypothesize that its small captive population may harbor TCC risk variants at the germline level. To aid conservation efforts and investigate the genetics of TCC, we present a new fishing cat chromosome-scale assembly, reaffirm its close genetic relationship with the Asian leopard cat (*Prionailurus bengalensis*), and identify and characterize single nucleotide variants (SNVs) from whole genome sequencing (WGS) data of healthy and TCC cats. Among genes previously associated with bladder cancer risk in human *BRCA2* was found to have the highest number of missense mutations in fishing cats, with only two variants exhibiting a predominance in TCC cats. These new fishing cat genomic resources will aid efforts to improve their genetic fitness and enhance the comparative study of feline genomes.

## Introduction

For many species, the creation of healthy zoo-managed populations is one way to promote longterm species survival, as the release of zoo-born mammals into the wild is not commonly practiced [1]. However, if a disease does occur in zoo-managed environments, it is of the utmost importance to determine its origin [2, 3]. Although habitat loss is the greatest threat to wildlife and natural ecosystems [4], studying the genetics of managed populations can provide insight into their wild ancestry, overall genetic fitness, and facilitate future disease investigations [2, 3].

The fishing cat (*Prionailurus viverrinus*) weighs between 11 and 35 pounds, inhabits the wetland habitats of Southeast Asia, and, unlike most other felines, relies heavily on waterways for food [5]. Although primarily piscivorous, these opportunistic nocturnal predators also feed on small mammals, amphibians, reptiles, crustaceans, and birds, albeit to a much lesser extent [6, 7]. The fishing cat has a stout, muscular body, an elongated head, webbed paws, and a shortened tail, all of which are well adapted for their aquatic lifestyle. Within felid phylogeny, it is part of the Leopard Cat lineage, which consists of the Asian leopard, flat-headed, rusty-spotted, fishing, and Pallas cats, and is flanked by the domestic and Bay cat lineage outgroups [8].

The Association of Zoos and Aquariums (AZA) has developed Species Survival Plans (SSP) for numerous at-risk species, including the fishing cat. The SSP helps coordinate breeding efforts to maximize genetic diversity from limited gene pools [2], but it is important to note that not all fishing cats were originally managed in AZA-accredited institutions and are therefore not included in the current fishing cat SSP. The historical fishing cat population in North American zoos can be traced back to founder individuals in the early 1900s, but the majority of founders were imported in the 1960s and 2000s from Sri Lanka, Thailand, and Cambodia. There are currently 25 captive-born individuals managed by the fishing cat SSP at various AZA-accredited institutions and housed in their own enclosures within small mammal houses.

Disease is closely monitored in all zoo-managed species for trends that could impact sustainability. The appearance of diseases with possible genetic causes or risk factors is especially concerning for smaller zoo-managed populations. It is therefore concerning that TCC accounted for 13% of all zoo-managed fishing cat deaths between the years of 1995 and 2004 [9]. Transitional cell carcinoma has been described in multiple species, including cattle, dogs, cats, some marine mammals, and humans [6, 9] yet its exact cause remains unknown. For dogs, TCC is the most common lower-urinary tract cancer; conversely, it is rare in domestic cats [10]. It also appears in older fishing cats, with the most common clinical sign being persistent hematuria (blood in the urine), and tumors occurring most commonly in the trigone region of the bladder [9]. This is consistent with tumor location in the domestic dog and cat [10]. For captive fishing cats, due to their higher relatedness relying on their current pedigree structure it is possible the genetic etiology of TCC is associated with the segregation of risk alleles [9]. Due to the known risk that deleterious alleles pose to small populations, an investigation into possible germline variants conferring risk is warranted. In humans, the higher occurrence of germline alleles in bladder cancer compared to healthy individuals was evaluated in a cohort of 1,038 patients that led to a total of 11 genes predicted to confer higher risk of bladder cancer emergence [11]. This offers a starting point of candidate genes worth examining in other species susceptible to TCC. However, for the fishing cat population, a few challenges with this approach include a small cohort size, availability of samples, age-related presentation, and the chance that fishing cat orthologs of human germline risk genes may not confer risk in TCC.

Highly contiguous genome assemblies, some with gap-free chromosomes, are now becoming readily available for some species [12, 13]. The currently available fishing cat assembly, PriViv1.0, is highly fragmented with 142,198 total contigs, none of which are assigned to chromosomes, presenting challenges for whole-genome alignment and variant-calling, especially of structural variants (SVs) [14]. We present a new high-quality fishing cat reference genome that is crucial for examining genome synteny among multiple felid species toward a better understanding of phenotypic differences among Felidae species and performing variant discovery for studying disease etiology. Our study focus was to detect putative risk alleles present in candidate genes for TCC that may allow for better genetic management of the fishing cat population. With this new catalog of discovered sequence variants, we enable future studies that attempt to understand this type of bladder cancer and its occurrence in other species.

## Results

### Pedigree construction

The historic population in North American zoos is comprised of a total of 161 cats (Fig. 1a). By reconstructing this pedigree, we could identify individuals that would be most informative for investigating germline alleles potentially associated with TCC occurrence. Over the past 15 years, 15 fishing cats were diagnosed with TCC, with blood or tissue samples from five TCC cats used in this genetic analysis. For all cats classified as TCC-affected, TCC was verified by zoo pathology reports and histologic confirmation of tumor within the bladder wall (Fig. 1b). Using the 2019 fishing cat studbook, each living cat was traced back over 12 generations to the founding individuals and their respective origins, primarily from the countries of Sri Lanka, Thailand, and Cambodia (Fig. 2a). Some founders were labeled as Asia origins but we cannot establish a country of origin. Although there are just 25 cats currently managed in AZA zoos, the historical SSP population numbered as many as 60 cats. From this cohort, we were able to identify a reference individual (Fig. 2b), as well as all cats selected for WGS and RNA sequencing (RNAseq) (Fig. 1a).

**Figure 1:**
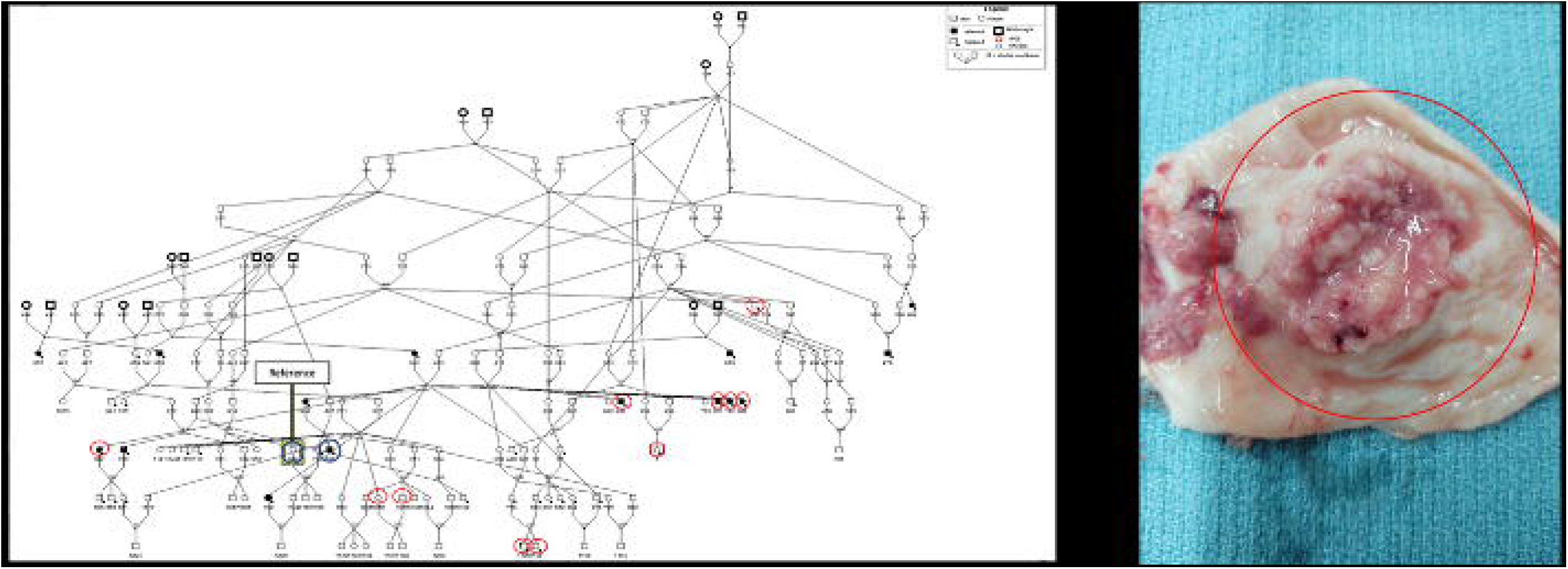
Studbook pedigree and transitional cell carcinoma (TCC) tumor. a. Depicted here is the North American fishing cat population pedigree. In total there are 161 cats in the historic population, with 25 of those being cats currently in zoo-managed care and 15 total cats being confirmed as TCC cases. Males are indicated by squares and females are circles. TCC cats are filled entities. Cats with a dot next to them are those in which samples (blood, urine, tissue) are collected. Founder individuals are the bolded entitles and all known cancer cats are indicated by filled entities. All WGS sequences are indicated in red, and the RNASeq cats are indicated by blue circles. Reference cat is indicated by yellow arrow. b. Indicated by the red circle is a tumor excised from the bladder wall of a transitional cell carcinoma affected fishing cat. Photograph provided by Dr. Leslie Lyons, University of Missouri.

**Figure 2:**
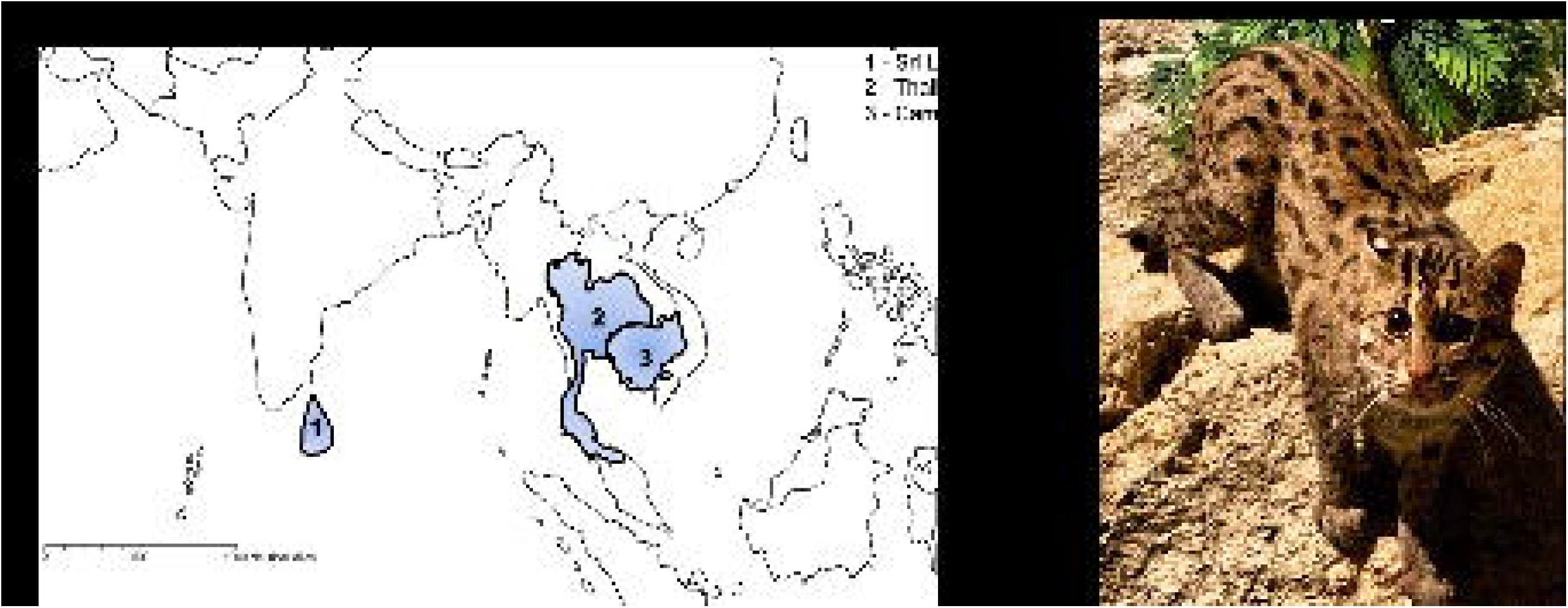
Geographic origins of founder individuals and Reference fishing cat “Anna”. a. Illustrated here are the geographical origins of the current North American zoo fishing cat population individuals according to studbook information. Of the 16 founder cats, 6 were from Sri Lanka (1), 5 from Thailand (2), and 2 from Cambodia (3). The remaining 3 were documented as being from Asia, however, no specific country of origin is indicated. b. Pictured here is the fishing cat selected for the reference genome assembly. Anna (AZA Studbook #950). Initially born in the La Fleche Zoo on 9/14/2010. After being transferred to two other European facilities in Bucharest, Romania, she eventually joined the AZA population in 2010 at the Chicago Zoological Society. She was then transferred to the Oklahoma City Zoo in 2020. Photograph provided by Animal Care Specialist Natalie Farley, Chicago Zoological Society.

The completed pedigree reveals multiple offspring resulting from consanguineous mating. Any consanguineous mating occurring between second cousins or closer were identified for the purposes of this study [15]. Some examples included in the Studbook (SB) are animal 226 mated with 229 (full sib to 226’s dam) to produce offspring 431,433, 37, 456, 306, 356 and a full-sib mating pair of 175 and 176 that produced 213, 212, 298 and 254 at the top right of the pedigree (Fig. 1a). Many of these closely-related individuals went on to produce at least one fishing cat in the pedigree. On the right side of the pedigree the founder cats 168 (Thailand), 183 (Thailand), and 170 (Sri Lanka) all were born between the 1970-80s and were initially bred and managed outside of North American zoos. Additional founders on the left side of the pedigree, including 491-494,182 (Sri Lanka), 503, 569-570 (Thailand), were all born in the 1980s and 1990s. In the early 2000s, two additional founders in the middle of the pedigree, 653-654 (Cambodia), were introduced. The number of consanguineous mating events fortunately occurred in earlier generations with later changes implemented by the AZA and SSP groups that reduced inbreeding.

### De novo assembly

Based on her position in the current AZA fishing cat pedigree and tissue availability, we chose Anna (AZA studbook #950, Fig. 2b), an 11-year-old fishing cat from the Oklahoma City Zoo, for long-read sequencing and genome assembly. At time of death Anna was determined to be TCC negative. We generated 30x sequence coverage of PacBio HiFi reads and assembled these into 245 primary contigs using HiFiasm [16]. Contig scaffolding was accomplished using ~35x sequence coverage of a Hi-C library [17, 18] that resulted in 19 chromosomes and 172 unplaced sequences (Supp. Fig. 1a and 1b). Manual visualization of MashMap [19] alignments against the most closely related species, Asian leopard, verified chromosome orientation (Supp. Fig. 2a, 2b). The total assembled size was 2.46 Gb with N50 contig and scaffold lengths of 68.7 Mb and 144.9 Mb, respectively, with 96.3% of sequence assigned to chromosomes. These assembly metrics were compared to the single haplotype assemblies of domestic cat and Asian leopard cat genomes, and despite the non-phased state of the fishing cat assembly was found to be comparable in assembly contiguity [13] (Table 1). In contrast, given the short-read composition of the prior fishing cat assembly (PriViv1.0) [14] our reference in comparison increased in size by nearly 16 Mb, and contiguity as measured by N50 contig length ~2,000 fold [14].

**Table 1:** NCBI statistics for select feline reference assemblies. Outlined here are the assembly statistics for each feline reference genome from the NCBI database. Although domestic cat and Asian leopard cat are all phased assemblies, the fishing cat quality statistics are within the same range, especially when comparing assembly sequence length, scaffold N50, and contig N50.

|  | Domestic cat<br>(F.catus_Fca126_mat1.0) | Asian leopard cat<br>(Fcat_Pben_1.1_paternal_pri) | Fishing cat (UM_Priviv_1.0) |
| --- | --- | --- | --- |
| Total sequence length | 2,425,747,038 | 2,435,718,761 | 2,461,562,329 |
| Total ungapped length | 2,425,739,938 | 2,435,706,361 | 2,461,531,252 |
| Number of scaffolds | 71 | 84 | 192 |
| Scaffold N50 | 148,491,486 | 148,587,958 | 144,906,162 |
| Number of contigs | 110 | 140 | 255 |
| Contig N50 | 90,731,473 | 82,622,880 | 68,766,634 |

### Genome annotation

Prior to gene annotation we performed Benchmarking Universal Single Copy Orthologs (BUSCO) analysis to assess genome completeness [20]. In the fishing cat genome, 93.5% of BUSCOs were complete, with only 4.7% missing compared to 5.0% in Asian leopard cat [13] (Supp. Table 1). In general, all BUSCO [20] findings were consistent with other highly-contiguous feline genome assemblies (Supp. Table 1). To validate *in silico* gene predictions, we generated RNAseq data for two fishing cat individuals Kiet (TCC positive at time of death) (SB #780) and Anna (TCC negative at time of death) (SB #950), from bladder and kidney tissue, respectively. Protein-coding and non-coding genes were predicted using the standardized NCBI RefSeq genome annotation processes [21] (see Supp. Table 2 for the annotation report). In sum, we find 20,055 protein-coding genes and 68,764 transcripts with 6,904 being non-coding genes. Our protein-coding gene total is consistent with predicted gene counts for other feline species [13] (Supp. Table 3).

### Structural analysis

Cross-species whole genome comparisons between recently diverged species can highlight signatures of evolutionary adaptation and speciation. We evaluated the sequence structural similarities and differences between the fishing cat and Asian leopard cat assemblies Fcat_Pben_1.1_paternal_pri [13], which diverged from fishing cat ~3 million years ago (Supp. Fig. 3). Three approaches were followed: Data-Visualization asm2ref pipeline [22] to generate chromosome synteny plots, SafFire [23] to evaluate order and orientation differences, and Assemblytics [24] to detect insertions, deletions, tandem expansions and contractions, and repeat expansions and contractions. Overall, manual reviews of whole genome alignments confirmed an expected high amount of one-to-one synteny (Fig. 3a). SafFire [23], but one obvious distinction was a large putative chrD1 inversion (6 Mb) (Fig. 3b). The chromatin proximity map shows the highest probability supporting its current inverted orientation and the inversion breakpoints reside in an un-gapped region that each suggest this inversion to be a natural difference to the Asian leopard cat. However, further confirmation in additional fishing cat genomes will be needed to be certain.

**Figure 3:**
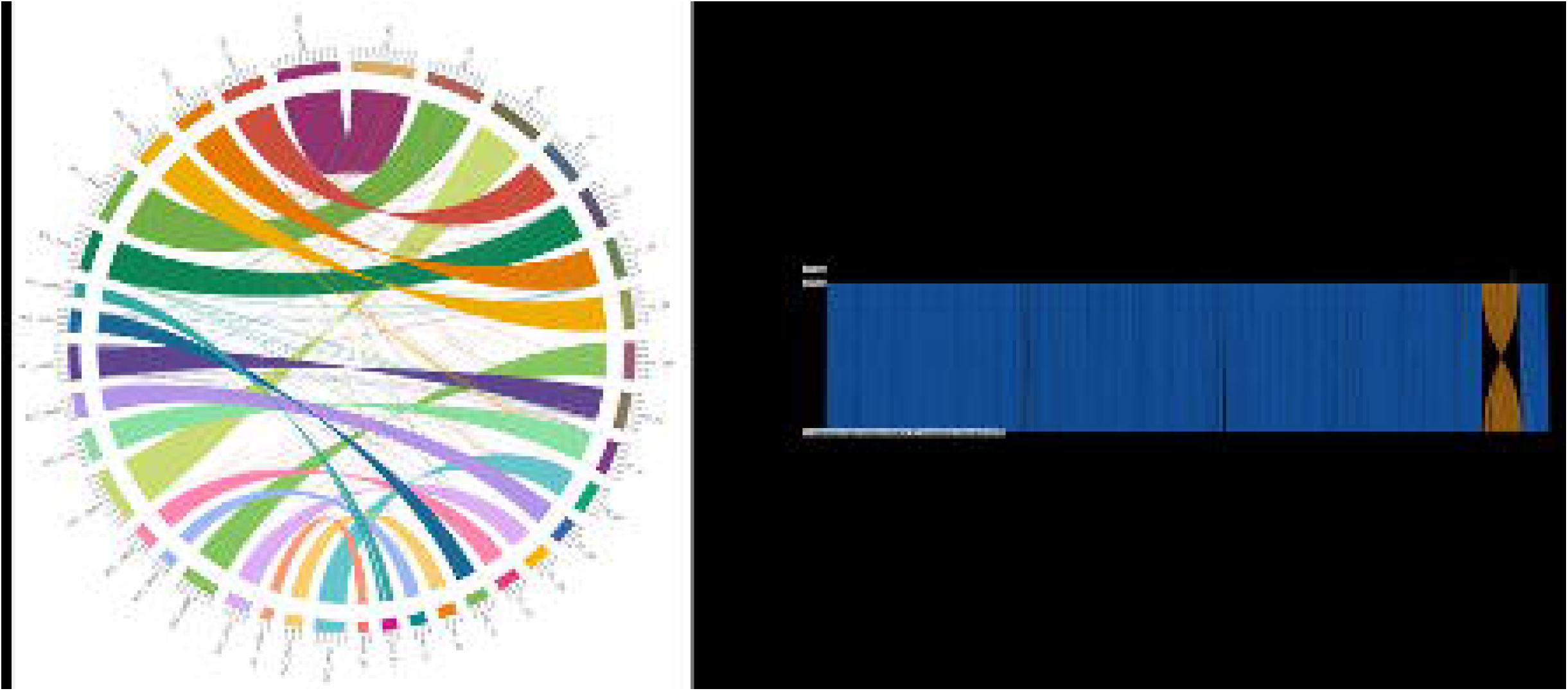
Chromosome alignments between the Asian leopard cat and fishing cat. a. Pictured is the circos plot comparing the Asian leopard cat chromosomes (ALC) to the fishing cat chromosomes (FC). Most alignments pictured are found to exhibit a high level of synteny with each other. b. One chromosome with an interesting alignment is in D1 as indicated by this SafFire output. D1 alignments illustrate an inversion ~6.0 Mb in size towards the end of the alignment.

We further refined our inter-species knowledge of major structural variation using AssemblyTics. In this output, we evaluated SVs and repeat variation that included estimates of tandem repeat expansions and contractions [24] (Supp. Table 4). In comparison to the Asian leopard cat, we find a total of 95,556 SVs with the most being 39,358 insertions. For repeats, 2,532 tandem expansions, 1,393 tandem contractions, 8,368 repeat expansions, and 15,626 repeat contractions were identified (Supp. Table 4).

### Cohort sequencing and variant annotation

In total, we generated whole genome sequencing (WGS) data on 11 cats (5 TCC and 6 unaffected) (Supp. Table 5) at ~25x sequence coverage on average to detect single nucleotide variants (SNVs) and indels. We then characterized the variant types with a focus on missense (variation causing local amino acid changes to coding sequence) and nonsense (predicted full length protein truncation) classifications (Supp. Table 6). Our cohort size was limited due to the availability or quality of samples. Initially, we used the Genome Analysis Tool Kit (GATK) [25] default parameters to detect a total of 7,541,694 SNVs and 2,600,943 indels. To reduce the presence of false positives we replotted SNVs and indels from GATK default parameterization [25] and chose the optimal inflection in the QD filter at <5 for both SNVs and indels and report a hard filtered total of 7,431,632 SNVs and 2,453,263 small indels ranging from 1-291bp in size. We also found 10,704 multiallelic SNV sites that were not considered in further analyses.

We used the Ensembl Variant Effect Prediction (VEP) [26] tool to categorize variants (Supp. Table 6), summarize their mutational load, and possible TCC impact. Our focus was on the interpretation of SNVs as this class of variants are less prone to error compared to indels. For all discovered variants, 74.4% were SNVs, 10.1% insertions, 9.4% deletions, and 6.1% sequence alterations (size of variant change is equal to that in the reference). For protein-coding consequences, most SNVs were synonymous at 57.5% (101,309 total variants), while 38.4% were missense (67,664 amino acid altering variants; Fig. 4a). The most severe in protein disruption, nonsense variants (stop loss, start loss, stop gained, and frameshift variants) were rare as expected at 4.1%. This included 2,587 frameshift, 547 stop gain, 101 stop lost, 146 start lost, 1,555 in frame insertions, 1,969 in frame deletions, and 54 protein altering variants (Supp. Table 6).

**Figure 4:**
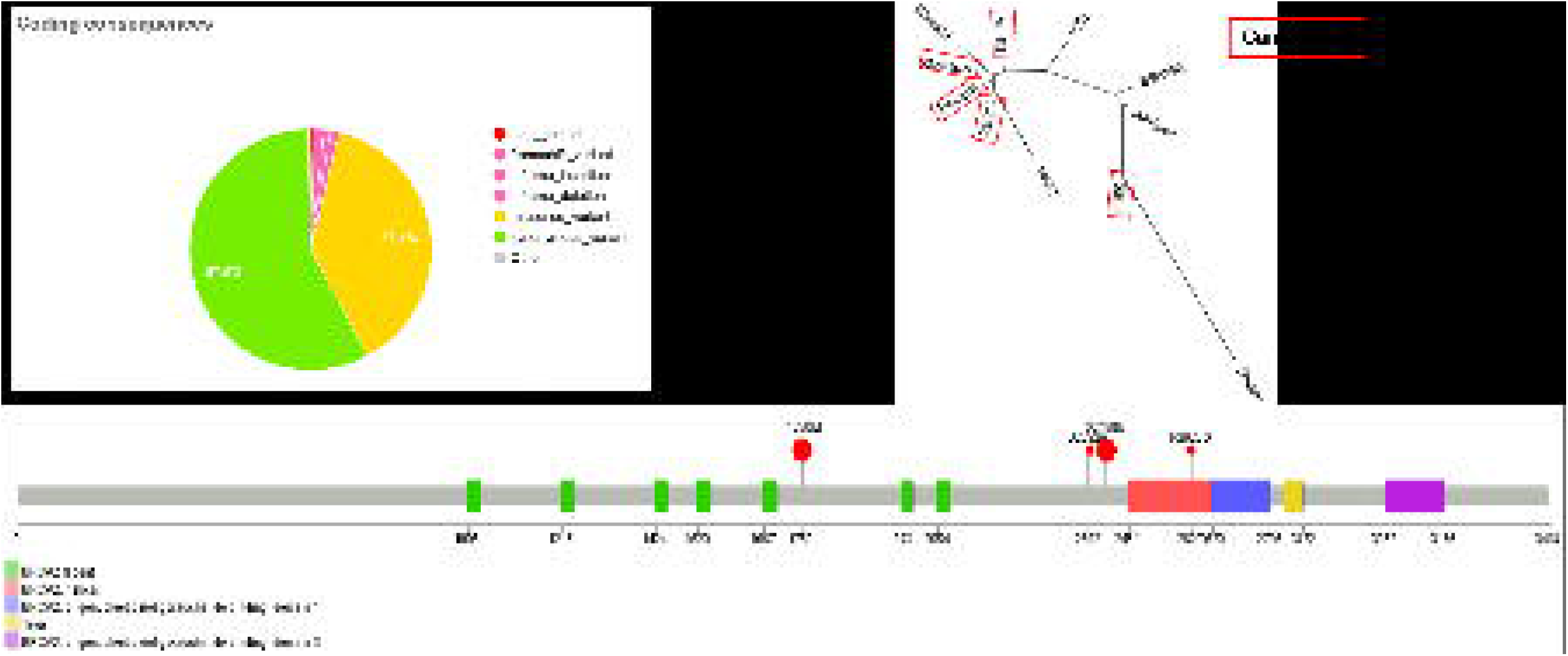
Missense variants identified in the fishing cat cohort. a. Here are the coding consequences of the variants identified in the fishing cat cohort. 57.5% were identified as synonymous, and 38.4% were identified as missense. Since missense variation is able to affect subsequent protein formation and function, any cancer genes of interest will be examined for missense variants. **b.** The relationship tree depicted here shows all cats evaluated in the fishing cat cohort. Individuals in red indicate cancer confirmed individuals. Sibling groups SB1195 and Juniper, as well as Gorton, Pavarti, Padma, and Maliha are grouped together. **c.** This lollipop plot illustrates 4 of the missense locations present in all cancer cats in *BRCA2* (fishing cat gene ID: LOC125162678). In each instance, all cancer cats expressed these missense mutations at these locations, with some normal-presenting cats also experiencing missense as well. All 4 sites experienced an amino acid change at each of these sites as a result of the base pair changes. T1750M and G2430E are indicated by larger points due to a higher 90.9% prevalence in the cat cohort. The smaller lollipops (D2392G and R2620G) indicate positions more strongly correlated with cancer cats, with 50% of non-cancer cats also affected.

### TCC candidate genes

Due to the lack of prior research into the genetics of TCC in fishing cats, we implemented a candidate gene strategy by searching for variants in the fishing cat orthologs of eight human genes associated with bladder cancer risk that includes *BRCA1, BRCA2, CHEK2, ATM, MSH2, MUTYH, MITF*, and *MLH1* [11]. These candidate genes considered in this study all have been previously defined as either playing roles in tumor suppression or DNA repair [27–29]. A total of four genes showed the presence of missense variants while none were seen in *MSH2, MUTYH, MITF*, or *MLH1* (Table 2). No nonsense variants (start codon, stop codon, splice donor, splice acceptor) were noted any of the evaluated genes. The *CHEK2* (68) had the highest followed by *BRCA2* (66), and *BRCA1* (33) genes (Table 2; Supp. Fig. 4). Among these three genes harboring missense variants, only one, *BRCA2*, had two independent variants that were found in all TCC cats (Fig. 4b, 4c). The two missense positions in *BRCA2* are protein positions D2392G and R2620G yet these were present in 50% of unaffected cats [30] (Fig. 4c). Two other sites, T1750M and G2430E, were both present in 100% of cancer cats and 83% in normal (Fig. 4c). The occurrence of these variants in normal cats even when found in all TCC cats, suggest more sequenced individuals and scrutiny of unaffected cats is needed to resolve this discrepancy. At this point we are not able to assign any of these candidate gene variants as high risk for TCC development. distinguish between these scenarios with the current data. Both *BRCA1* and *BRCA2* had nonsense variants detected with *BRCA1* having two in frame insertions and *BRCA2* having six in frame deletions. However, their frequency among all our studied cats suggested unlikely association with TCC risk.

**Table 2:**
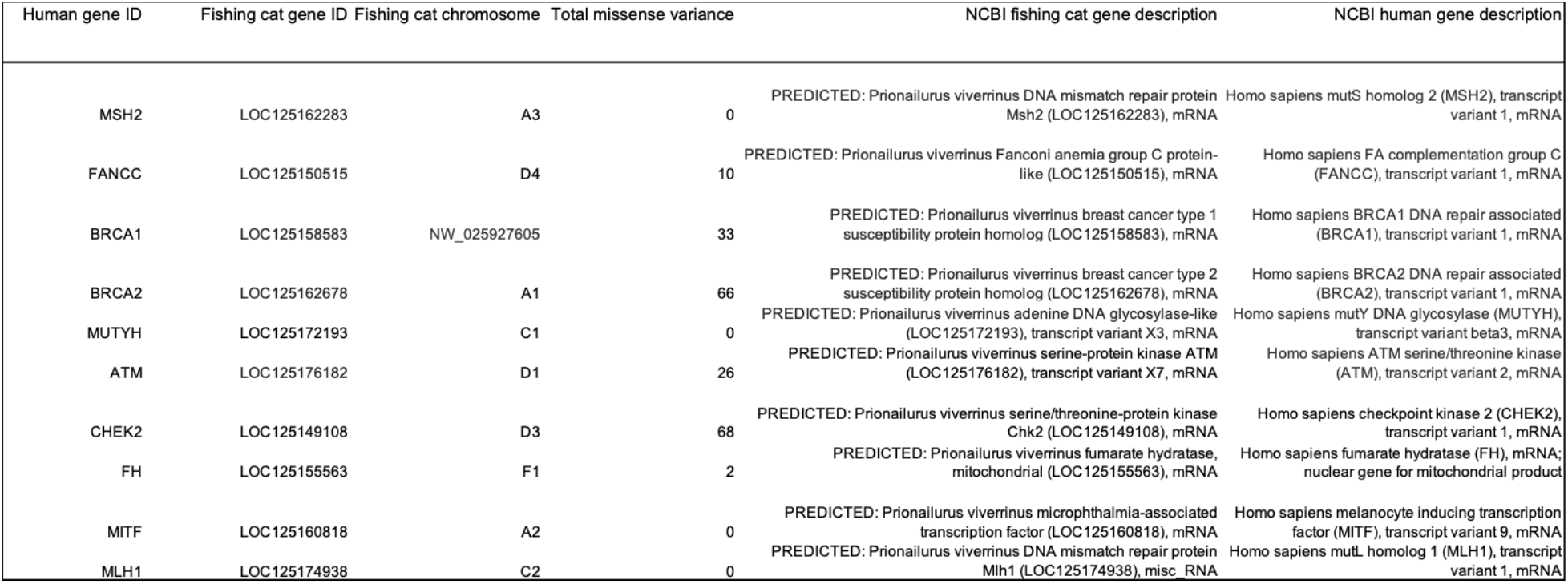
Pathogenic risk genes and associated missense variants per gene. Indicated are the human gene ID, homologous fishing cat gene ID, chromosome location on the fishing cat genome, total missense identified, and the description of each gene for the fishing cat and human as defined by NCBI Gene [31]. After blasting the genes against the fishing cat genome, the homologous gene IDs were evaluated in the VEP output for any missense variation happening within the fishing cat population. In total, the genes with the most missense variation present were CHEK2 with 68 variants, BRCA2 with 66 variants, BRCA1 with 33 variants, and FANCC with 10 variants. Gene descriptors for both fishing cat and human were obtained from the NCBI database.

## Discussion

In the wild, natural selection frequently eliminates disease risk alleles that reduce fitness, but not always. For diseases that manifest after reproductive age, such as cancer, these deleterious alleles continue to be transmitted to succeeding generations. In efforts to conserve species, captured population founders may harbor unknown germline disease risk alleles, which are subsequently propagated in controlled breeding schemas. If this were to occur, the availability of accurate pedigrees of captive populations would be crucial for understanding disease incidence and, if possible, determining its cause. Our construction of the largest fishing cat pedigree (n=161) with correct placement of all individuals is the first step in investigating the increased incidence of TCC observed in captive fishing cats, as only 25 captive-bred individuals remain in zoos [6, 9]. A pedigreed female was used to generate a chromosome-scale reference with a 2,000-fold improvement in assembly continuity compared to the previous scaffold-level assembly, Priviv1.0 [14]. In addition, the new fishing cat assembly quality metrics were comparable to domestic and Asian leopard cats, even with these genomes both the result of trio-binning approach, which allows for the phasing of single haplotypes compared to our fishing cat diploid assembly starting point [13]. Sequence completeness and accuracy demonstrate that this new fishing cat reference is the optimal computational resource for investigating fishing cat diseases, its population diversity, and Felidae interspecies genome evolution.

Various measures of gross co-linearity indicate that Felidae genomes are highly conserved for most chromosomes **[32, 33]**. Within the felid phylogeny, the fishing cat is the sister lineage to the Asian leopard cat **[8]**, which prompted us to search for any major structural changes that have occurred within the past 3Myr since their divergence. We confirmed substantial conserved synteny in the *Prionailurus* genus utilizing a variety of alignment techniques. Given the high level of genome-wide synteny at the moderate and deepest divergence of the cat family in comparative studies of the domestic cat to the Asian leopard cat and lion **[13, 34]**, this result is not unexpected. Nonetheless, interesting small scale and rarely large deviations in sequence order and orientation can arise even in short spans of evolution. We find a 6Mb inversion on chromosome D1 between these two *Prionailurus* genomes. Other differences, such as those at the ends of chromosomes D4 and E2, suggest assembly artifacts (Supp. Fig. 5a, 5b) and collectively all interspecies SV differences will require additional experimentation within species. Independent pairwise alignments of the assembled contigs between these *Prionailurus* species reveal numerous varied size insertions, deletions, and repeat expansions and contractions. Over the relatively evolutionary-short *Prionailurus* period of species divergence, fishing cat tandem repeat expansions accounted for the greatest totals by category, followed by contractions. However, across felid species repeat expansions did not drive their expansion to genome size equivalency with other mammalian species. Genomes of fishing cats and other felid species, have 34% total interspersed repeats compared to ~50% in most other mammalian genomes **[21, 35, 36]**. This fits the observation that carnivore genome size is on average ~400Mb smaller and suggests repeats have shaped the carnivore genome structure very differently in the mammalian eutherian lineage.

The possibility that disease risk alleles even though rare in wild populations can be unknowingly segregating in small zoo populations, started from wild individuals, is often not discovered until pedigrees are well established. The clinical symptoms of possible bladder cancer were first observed as far back as 1991 and later proposed to be TCC in captive fishing cats **[6]**. Although genetic cause, including no proposed mode of inheritance, was found its occurrence throughout the pedigree warranted further genetic evaluation. Our study provides the first estimation of segregating SNVs and indels in fishing cats and their use investigating disease. Of the missense variants we examined in fishing cat gene orthologs for human bladder cancer risk genes, only *BRCA2* on chromosome A1 showed a skewed distribution in TCC over healthy cats. In addition, one control cat, Wasabi, shared missense variants at all four *BRCA2* sites, as well as the same missense genotype (all cats heterozygous from the reference) as all the cancer cats at positions D2392G and R2620. However, despite this shared genotype, there are no clinical signs of TCC presented in this cat. Given the close phylogenetic relationship between Wasabi and the sibling group of cancer cats (Padma, Pavarti, Gorton, Maliha) (Fig. 4b), it is probable that this cat carries the same risk allele as the cancer group if germline inheritance is in fact promoting TCC. However, due to its young age, this cat has not yet displayed clinical signs of TCC; therefore, future monitoring of this cat and its siblings (as shown in Fig. 1a) may be necessary. In humans, *BRCA2* alleles are well-documented for the higher occurrence of familial breast cancer and ovarian cancer, as well as pancreatic and prostate cancer **[28, 37, 38]** but historically, *BRCA2* has had no significant association with any bladder cancer occurrence. Still, recent publications are now considering a possible *BRCA2* role in some types of human bladder cancer predisposition **[11, 28, 39]**. Furthermore, it’s function in DNA repair and homologous recombination (HR), with acquired, mutations triggering the malfunction of other DNA damage repair mechanisms and the loss of tumor suppression indicate importance in TCC **[38]**. The finding of increased numbers of germline variants associated with DNA damage repair pathways in human urothelial carcinoma (UC) patients (such as *BRCA1/2. CHEK2, ATM*), particularly in *BRCA2* **[11, 39]** is again interesting for TCC. In fact, one group has called for *BRCA2* future germline UC screenings **[39]**. A recommendation we can’t make at this time due the small scope of our TCC sampling, younger age at sample collection, and some cats which may harbor risk alleles but show no clinical symptoms.

As sequencing technology is very affordable today, examining a larger cohort of TCC-susceptible species will help clarify causative risk variants in all. Moreover, this preliminary fishing cat research highlights the importance of screening small captive populations for known disease risk genes in other species, particularly using the depth of knowledge available in human, to guide mating decisions and more importantly warning their caretakers of future disease occurrence if the predictive data allows. We propose these genome-wide findings illuminate their genetic fitness, with a goal to diminish the occurrence of TCC in this fragile population, and thus promoting future fishing cat conservation efforts.

## Methods

### Pedigree construction

A pedigree was first drawn using the program Pedigree-Draw to select the reference individual that accurately represents the Association of Zoos and Aquariums (AZA) population. Using the current fishing cat studbook from 2019, each living cat was traced back over 12 generations to the founding individuals and their respective origins.

### Reference individual

Anna, an 11-year-old female fishing cat from the Oklahoma City Zoo, was selected as the candidate for genome assembly. Through pedigree assessment, she was found to be an individual that accurately represented the fishing cat species and the AZA zoo-managed population. High molecular weight (HMW) DNA was obtained from frozen kidney tissue extracted during necropsy and stored at −80C. The HMW DNA was isolated using the 10x Genomics Demonstrated Protocol: DNA Extraction from Single Insects (10x Genomics). The final HMW DNA quantity was determined using the high-sensitivity Invitrogen Qubit Fluorometer protocol. Final HMW quality was determined using a 0.7% agarose gel and imaged on the Uvitec Cambridge Uvidoc HD6 UV Fluorescence and Colorimetry instrument.

### Genome long-read sequencing

HMW DNA was used for library construction and sequencing on a Sequel II instrument using HiFi mode (PacBio). A target of ~20 kb fragment size was used to construct SMRTbell libraries using CCS Express Library Kit V2. Nine ug of starting DNA was used with small fragment removal before shearing, size using a megaruptor shear speed 31, and sized on a Blue Pippin Instrument 10-50 kb. The final library concentration was 38.6 ng/ul in 15 ul, and a final size of 18 kb was used. In total three SMRT cells were generated to an estimated coverage of HiFi reads at 30x.

### Assembly construction and curation

De novo assembly used HiFi processed reads >18kb and was performed with Hifiasm (version 0.13-2208) [16]. BUSCO (version 4.1.2_cv1) [20] analysis with the arguments “-m genome –l mammalia_odb10” were used to evaluate the completeness of the genome [13]. We used the purge_haplotigs pipeline [40] to generate a cutoffs histogram. No haplotig curve was observed on the histogram so cutoff parameters were not utilized [41]. After running the purge haplotigs pipeline a re-run of BUSCO [20] confirmed estimated gene duplication rate. The remaining contigs served as input for scaffolding using multiple programs. Hi-C data generated by the DNA Zoo [17, 18] was used to produce scaffold structure using the hic-saffolding-nf pipeline [42] with Salsa (version 2.2) and Juicebox (version 1.11.08) [43] to evaluate the resulting Hi-C heat maps for order and orientation convergence.

Misplaced scaffolds were identified using multiple programs and orthogonal evidence to estimate the most accurate chromosome order and orientation. Chromosome nomenclature was used in accordance with the original genetic linkage map groupings [44]. MashMap (version 2.0) [19] was used to compare the fishing cat scaffold assembly to the current domestic cat genome assembly Felis_catus_9.0 [45] to further correct assembly error. Using Juicebox [43] and MashMap [19] together, Agptools [46] was used to adjust or move contigs to the predicted correct placement that was accomplished by flipping, breaking, and joining contigs together. Chromosome representation and accuracy benchmarking analysis identified 37 scaffolds requiring error correction, such as chrB2. Evaluating the MashMap [19] output of the fishing cat assembly aligned to the domestic cat [45] (Supp. Fig. 2a) revealed fishing cat scaffold_4 covered the entirety of domestic cat chrB2, yet it was necessary to break this scaffold into three pieces and join them back together for correct order and orientation where accuracy was reconfirmed in the MashMap [19] and the Hi-C contact map. Importantly, if local order within a contig was not conserved, we avoided homogenizing the fishing cat assembly to mirror the domestic cat thus preserving the original fishing cat genome structure whenever orthogonal evidence supported it (Supp. Fig. 2b).Agptools was used to rename scaffolds to be consistent with the linkage groups of the domestic cat genome[44] and finally MashMap[19] was run again to verify the new assembly output corrected all detectable errors.

### RNA sequencing and gene annotation

Total RNA from the bladder and spleen of two fishing cats was isolated via the RNeasy Plus Universal Mini Kit (Qiagen). The bladder tissue sample was from a 12-year-old male “Kiet” from the San Francisco Zoo, and the spleen sample originated from an 11-year-old female fishing cat “Anna” from the Oklahoma City Zoo. The cDNA libraries of each sample were generated and sequenced on an Illumina Novaseq 6000 to coverage of ~50 million reads/library. Both RNA datasets were submitted to NCBI for annotation using the RefSeq annotation pipeline [21].

### Structural analysis

Three independent approaches were used in this analysis. AssemblyTics [24] was used to compare the fishing cat genome to its closest living relative, the assembled Asian leopard cat (Fcat_Pben_1.1_paternal_pri) [13]. Within AssemblyTics [24] we ran the MUMmer nucmer algorithm to align the fishing cat contigs as the query sequence, against the reference contig sequences of the Asian leopard cat. In our study, we used the default unique anchoring size of 10,000 bp [24]. The resulting delta file was input into the Assemblytics website and run on the default parameters: a unique contig sequence anchoring size of 10,000 bp, and the variants themselves classified by type and size ranging from 50-500 bp and 500-10,000 bp [24].

Data-Visualization [22] was utilized for the synteny analysis between UM_Priviv_1.0 and the Asian leopard cat assembly Fcat_Pben_1.1_paternal_pri [13]. For optimal performance, both assemblies had to have the same number of chromosomes with the same nomenclature. Because of this, all unplaced scaffolds from both assemblies were removed, and all chromosomes were renamed from NCBI format to the corresponding chromosome naming format. Minimap2 (version 2.24-r1122) [47] was used as the genome alignment program using asm20 for cross-species analysis. Once the alignment file was generated it was then used as input into the Data-Visualization asm2ref pipeline to generate the resulting circos plot.

SafFire (version 0.2) [23] was utilized for the synteny analysis across all chromosomes between UM_Priviv_1.0 and Fcat_Pben_1.1_paternal_pri [13]. All unplaced scaffolds were removed, and chromosomes were renamed to optimize accurate alignment outputs between the two assemblies. Minimap2 (version 2.24-r1122) [47] was used as the genome alignment program using asm20 for crossspecies analysis. Rustybam (v0.1.30) [48] was then used to generate the input file for SafFire.

### TCC sample preparation and sequencing

A total of 11 cats with and without presenting TCC were selected for WGS and variant analysis. DNA was isolated from all samples using whole blood and tissue samples with the Qiagen DNeasy Blood and Tissue Kit (Qiagen). The DNA quantity and quality was determined using the Qubit Fluorometer instrument protocol (Invitrogen), and electrophoresis was performed on a 0.7% agarose gel with gel imaging on the Uvitec Cambridge Uvidoc HD6 UV Fluorescence and Colorimetry instrument, respectively. All samples were sequenced on the Illumina Novaseq 6000 (Illumina) with coverage of ~20x per cat with cDNA libraries generated using the Illumina DNA Prep protocol (Illumina) using double bead selection to select larger insert sizes (~550 bp).

### Variant call pipeline

The Genome Analysis Toolkit (GATK) (version 4.1.8.1) [25] was used for variant identification in a population cohort of 11 fishing cats, both with and without presenting TCC (Supplementary table 4). The program was run using default parameters and HaplotypeCaller [49] allowing for the joint genotyping of germline variants in all sampled individuals. The default GATK hard filtering parameters for SNPs (QD<2.0, QUAL<30, SOR>3.0, FS>60.0, MQ<40.0, MQRankSum< −12.5, ReadPosRankSum < −8.0) and indels (QD<2.0, QUAL<30.0, FS>200.0, ReadPosRankSum < −20.0) were utilized for variant filtration after adjustments to the QualByDepth(QD) parameter threshold. To do this, filter plots to visualize the distribution of SNVs and Indels for our cohort were generated in RStudio (version 2021.09.0+351) using gridExtra and ggplot2. QD was adjusted (QD<5.0) for both SNPs and Indels as opposed to < 2 per best practice default. The remaining filters are left the same as QD accounts for most noise in the variant data. The Nextflow variant call pipeline was then re-run with the new filters and all statistics for the GATK VCF output were obtained using BCFTools stats (version 1.14) [50].

### Neighbor joining trees

The fishing cat variant call format (VCF) file was converted to phlyip format using Vcf2phylip (version 2.0) [51]. Phylogenetic analysis of the fishing cat was conducted using MEGA (version 11) [52] through neighbor joining methods. Finally, a bootstrap analysis was run with 1000 replicates to evaluate branch correctness, and the Kimura 2 Parameter was used to estimate genetic distances between samples in each tree.

### Variant effect, prediction, and annotation

Ensembl’s Variant Effect Predictor (VEP) (version 101.0) [26] was used for this analysis. [26]This study used the downloadable VEP Perl script and the fishing cat NCBI RefSeq GFF file [21] as a predicted Ensembl gene set is not available.

### TCC candidate risk gene investigation

The genes identified by Nassar et al. in human publication were used as a reference here [11]. Each gene on this list was evaluated for homologous sequences in the fishing cat genome. This was done using the NCBI BLAST program [53]. Sequences found to have an e value of 0 were selected due to significant sequence homology. Once the homologous sequences were established, the Integrative Genomics Viewer (IGV) (version 2.13.2) [54] was used to identify any variants found within the selected genomic regions. This was done by uploading the reference genome and index, GFF file, and VCF file with all the population data to IGV [54]. The VEP [26] output for the fishing cat was also used to verify if these sequences were experiencing missense variation, which could potentially lead to protein alterations that further promote disease-causing genes. *BRCA2* lollipop plot was illustrated using Lollipops (version 1.5.2) [30].

## Acknowledgements

We thank the following North American zoo institutions that provided sample collection and shipment: Oklahoma City Zoo and Botanical Garden, Cincinnati Zoo and Botanical Garden San Antonio Zoo, Memphis Zoological Garden and Aquarium, San Francisco Zoo, San Diego Zoo, Point Defiance Zoo and Aquarium, Mill Mountain Zoo, Smithsonian National Zoo, and The Jackson Zoo. We also want to thank the Chicago Zoological Society fishing cat Animal Care Specialists for providing insight into fishing cat management and for providing a photograph of the reference cat “Anna”. Finally, the funding for all WGS data used for the population analysis and TCC investigation were covered under the University of Missouri Genomics Technology Core Tier 1 Sequencing Funds, with additional support provided by the Basis Foundation. WJM acknowledges support from National Science Foundation grant DEB-1753760. The computation for this work was performed on the high performance computing infrastructure provided by Research Support Solutions and in part by the National Science Foundation under grant number CNS-1429294 at the University of Missouri, Columbia MO.

DOI: https://doi.org/10.32469/10355/69802.

## Statistics and reproducibility

Assembly statistics for each reference genome were obtained from the NCBI website. BUSCO statistics for the fishing cat reference was determined using BUSCO (version 4.1.2_cv1) [20]. All SNP and Indel statistics from the variant calling pipeline were obtained using BCFTools stats (version 1.14) [50]. All sample requirements for DNA or RNA isolation were followed in accordance with the following protocols: 10x Genomics Demonstrated Protocol: DNA Extraction from Single Insects (10x Genomics), Qiagen RNeasy Plus Universal Mini Kit (Qiagen), and Qiagen DNeasy Blood and Tissue Kit (Qiagen).

## Ethics approval and consent to participate

This study was conducted in accordance with the Association of Zoos and Aquariums fishing cat Species Survival Plan coordinator Tyler Boyd. All samples collected were from AZA Accredited facilities and through the Feline Genetics and Comparative Medicine Laboratory at the University of Missouri.

## Data availability

All raw and processed data for this study are available at NCBI BioProject under accession PRJNA815338.

## Code availability

Scripts used for this study are available at the following GitHub repositories: https://github.com/WarrenLab/hic-scaffolding-nf https://github.com/WarrenLab/purge-haplotigs-nf https://github.com/WarrenLab/agptools

## Author contributions

WCW oversaw the experimental design of the study. LAL conceived the study and provided samples for WGS DNA extraction. WJM provided other cat assemblies for comparison and consultation on feline assembly comparisons. RAC coordinated sample collection and performed the DNA and RNA isolations. RAC and ESR performed the genome assembly. RAC and LC performed variant calling. RAC performed all other computational analyses and interpreted results. KAT and WFS provided necessary insight into TCC occurrence in the fishing cat population. TB provided all Studbook information for the pedigree analysis and WGS sample selection. RAC wrote the manuscript with input from all authors.

