## supplementary tables for "A novel fishing cat reference genome for the evaluation of potential germline risk variants"

| | Domestic cat (F.catus_Fca126_mat1.0) | Asian leopard cat (Fcat_Pben_1.1_paternal_pri) | Fishing cat (UM_Priviv_1.0) |
|---|---|---|---|
| Complete | 8,616 | 8,610 | 8,623 |
| Percent Complete | 93.39% | 93.32% | 93.50% |
| Single-copy | 8,589 | 8,589 | 8,599 |
| Duplicated | 27 | 21 | 24 |
| Fragmented | 156 | 153 | 167 |
| Missing | 454 | 463 | 436 |
| Percent Present (Comp + Frag) | 95.08% | 94.98% | 95.30% |

**Supplementary Table 1: BUSCO scores. Here are the BUSCO scores for the selected feline assemblies.** Using the same BUSCO mammalia_odb10 dataset, the final scores for completeness, duplication, fragmented, and missing are all consistent with each other.

| RefSeq Annotation Report | | | | | |
|---|---|---|---|---|---|
| Feature | Count | Mean length (bp) | Median length (bp) | Min length (bp) | Max length (bp) |
| Genes | 26,992 | 44,238 | 13,021 | 49 | 2,066,153 |
| All transcripts | 68,764 | 3,182 | 2,579 | 49 | 103,095 |
| mRNA | 56,277 | 3,505 | 2,861 | 147 | 103,095 |
| misc_RNA | 3,687 | 3,283 | 2,801 | 117 | 15,837 |
| tRNA | 751 | 76 | 73 | 59 | 91 |
| lncRNA | 6,003 | 1,493 | 896 | 97 | 20,187 |
| snoRNA | 589 | 111 | 105 | 49 | 329 |
| snRNA | 1,113 | 114 | 107 | 60 | 199 |
| rRNA | 311 | 905 | 119 | 119 | 4,645 |
| Single-exon transcripts | 1,927 | 1,188 | 948 | 147 | 14,422 |
| coding transcripts (NM_/XM_ ) | 1,927 | 1,188 | 948 | 147 | 14,422 |
| CDSs | 56,290 | 2,049 | 1,509 | 96 | 103,095 |
| Exons | 264,672 | 306 | 137 | 1 | 22,878 |
| in coding transcripts (NM_/XM_ ) | 242,283 | 296 | 136 | 1 | 22,878 |
| in non-coding transcripts (NR_/XR_ ) | 44,225 | 305 | 135 | 2 | 18,596 |
| Introns | 236,897 | 6,513 | 1,562 | 30 | 937,235 |
| in coding transcripts (NM_/XM_ ) | 220,721 | 6,370 | 1,528 | 30 | 937,235 |
| in non-coding transcripts (NR_/XR_ ) | 37,542 | 6,607 | 1,761 | 30 | 861,863 |
| | Mean | Median | Min | Max | |
| Number of transcripts per gene | 2.59 | 1 | 1 | 50 | |
| Number of exons per transcript | 11.87 | 9 | 1 | 312 | |

**Supplementary Table 2: NCBI RefSeq annotation summary.**
Detailed in this table are the statistics for the NCBI RefSeq output for NCBI Prionailurus viverrinus Annotation Release 100. This table also shows the number of transcripts per gene and exons per transcript in the fishing cat gene annotation.

| Species | Total Protein Coding |
|---|---|
| Fishing cat (UM_Priviv_1.0) | 20,055 |
| Asian leopard cat (Fcat_Pben_1.1_paternal_pri) | 20,003 |
| Domestic cat (F.catus_Fca126_mat1.0) | 20,453 |

**Supplementary Table 3: Total Protein Coding Genes Across Feline Species.** Compared to the Asian leopard cat and Domestic cat phased genome assemblies the number of protein coding genes remains consistent across all 3 species.

| Structural Variant Type | Fishing Cat vs Asian Leopard Cat | | |
|---|---|---|---|
| | Size | Count | Total bp |
| Insertion | 50-500 bp: | 37142 | 8204841 |
| | 500-10,000 bp: | 2216 | 6140544 |
| | Total: | 39358 | 14345385 |
| | Size | Count | Total bp |
| Deletion | 50-500 bp: | 26655 | 5580183 |
| | 500-10,000 bp: | 1624 | 4851702 |
| | Total: | 28279 | 10431885 |
| | Size | Count | Total bp |
| Tandem Expansion | 50-500 bp: | 1991 | 470792 |
| | 500-10,000 bp: | 541 | 822596 |
| | Total: | 2532 | 1293388 |
| | Size | Count | Total bp |
| Tandem Contraction | 50-500 bp: | 1151 | 263668 |
| | 500-10,000 bp: | 242 | 398212 |
| | Total: | 1393 | 661880 |
| | Size | Count | Total bp |
| Repeat Expansion | 50-500 bp: | 5081 | 1321120 |
| | 500-10,000 bp: | 3287 | 10569301 |
| | Total: | 8368 | 11890421 |
| | Size | Count | Total bp |
| Repeat Contraction | 50-500 bp: | 9636 | 3031296 |
| | 500-10,000 bp: | 5990 | 13116375 |
| | Total: | 15626 | 16147671 |

**Supplementary Table 4: Genome structural comparisons.** This table summarizes the AssembyTics output comparing the fishing cat reference genome to Asian leopard cat reference. Overall, least differences are detected in tandem expansions (2,532 identified) and tandem contractions (1,393 identified).

| Studbook ID | Name | Sex | DOB | Age (years) | Sequencing | TCC status |
|---|---|---|---|---|---|---|
| 687 | Pavarti | F | 5/2/05 | 13 | WGS | Affected |
| 722 | Maliha | F | 2/23/06 | 13 | WGS | Affected |
| 693 | Sushi | M | 7/15/05 | 11 | WGS | Affected |
| 688 | Padma | F | 5/2/05 | 12 | WGS | Affected |
| 721 | Gorton | M | 2/23/06 | 11 | WGS | Affected |
| 1196 | Juniper | F | 3/10/16 | 6 | WGS | Normal |
| 356 | Fritz | M | 8/15/94 | 10 | WGS | Normal |
| 5 | Splash | F | 1/14/96 | 11 | WGS | Normal |
| 1095 | Wasabi | M | 5/17/13 | 9 | WGS | Normal |
| 1195 | SB #1195 | M | 3/10/16 | 6 | WGS | Normal |
| 1059 | Jonas | F | 7/31/12 | 10 | WGS | Normal |
| 780 | Kiet | M | 6/29/09 | 12 | RNAseq | Affected |
| 950 | Anna | F | 9/14/10 | 11 | RNAseq | Normal |

**Supplementary Table 5: Fishing cat sequencing cohort.** Listed in this table are the WGS cats and RNAseq cats, along with their studbook ID, name, sex, date of birth (DOB), current age, and TCC status. It is important to note that samples from normal presenting cats (except for studbook #356 and studbook #5 - both confirmed as non-TCC cats) were collected prior to 10 years old, when clinical signs of transitional cell carcinoma tend to occur. Studbook #950 is also the cat selected as the reference individual.

| Consequence type | Count |
|---|---|
| transcript_ablation | 2 |
| splice_donor_variant | 554 |
| splice_acceptor_variant | 555 |
| stop_gained | 547 |
| frameshift_variant | 2,587 |
| stop_lost | 101 |
| start_lost | 146 |
| inframe_insertion | 1,555 |
| inframe_deletion | 1,969 |
| missense_variant | 67,664 |
| protein_altering_variant | 54 |
| splice_region_variant | 30,675 |
| start_retained_variant | 9 |
| synonymous_variant | 101,309 |
| stop_retained_variant | 59 |
| coding_sequence_variant | 171 |
| 5_prime_UTR_variant | 61,396 |
| 3_prime_UTR_variant | 228,124 |
| non_coding_transcript_exon_variant | 69,156 |
| intron_variant | 18,048,749 |
| non_coding_transcript_variant | 1,695,493 |
| upstream_gene_variant | 1,517,673 |
| downstream_gene_variant | 1,508,744 |
| intergenic_variant | 5,224,881 |

**Supplementary Table 6: Fishing cat VEP output.** Detailed here are all the consequence types identified in the fishing cat cohort from Ensembl's VEP program. Most consequence types are found to be intron variants (63.2%) followed by intergenic variants (18.3%). In addition, a total of 67,664 missense variants were identified (0.2%).
