## supplementary figures for "A novel fishing cat reference genome for the evaluation of potential germline risk variants"

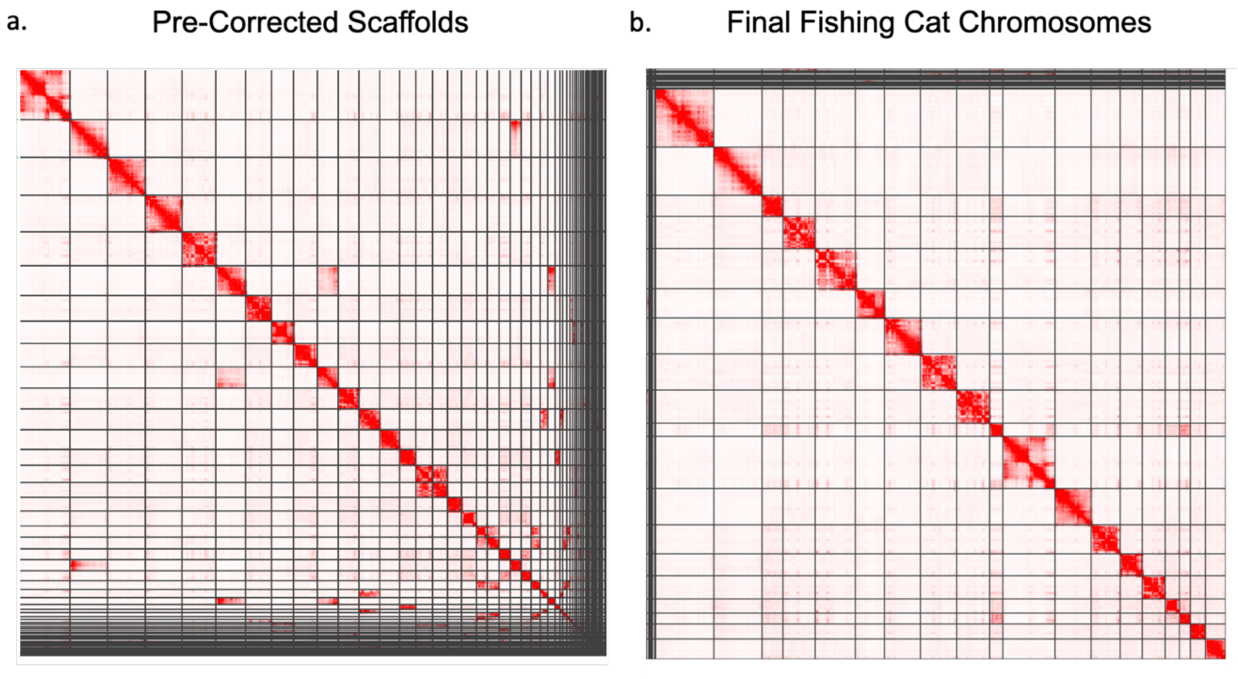


**Supplementary Figure. 1: Fishing cat Hi-C mapping outputs.**

a. Depicted here is the Hi-C image of the pre-corrected fishing cat scaffold level assembly.

b. The final Hi-C image of the 19 fishing cat chromosomes.

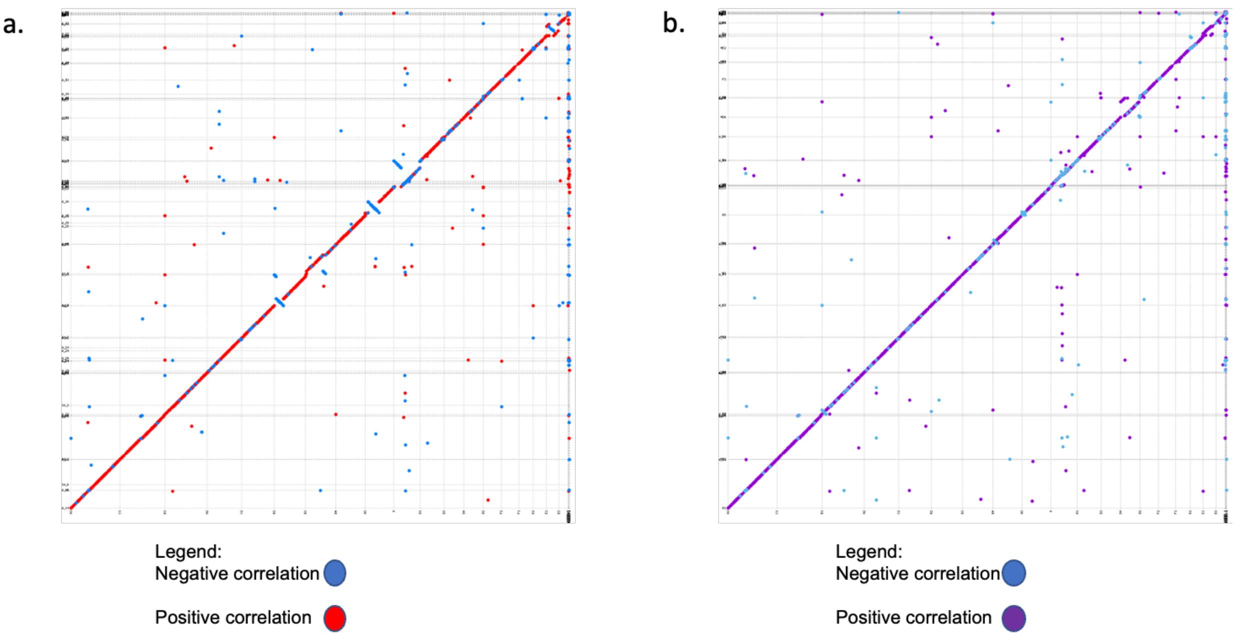


**Supplementary Figure 2: Scaffold correction using alignments to a similar feline genome.**

a. Pre-corrected MashMap dot plot comparing the fishing cat scaffold assembly (y) to the domestic cat assembly (x). Negative alignment correlations are blue and positive alignment correlations are red.

b. Finalized MashMap output of the fishing cat assembly (y) compared to the domestic cat assembly (x). Negative correlation alignments are indicated in blue and positive alignment correlations are purple.

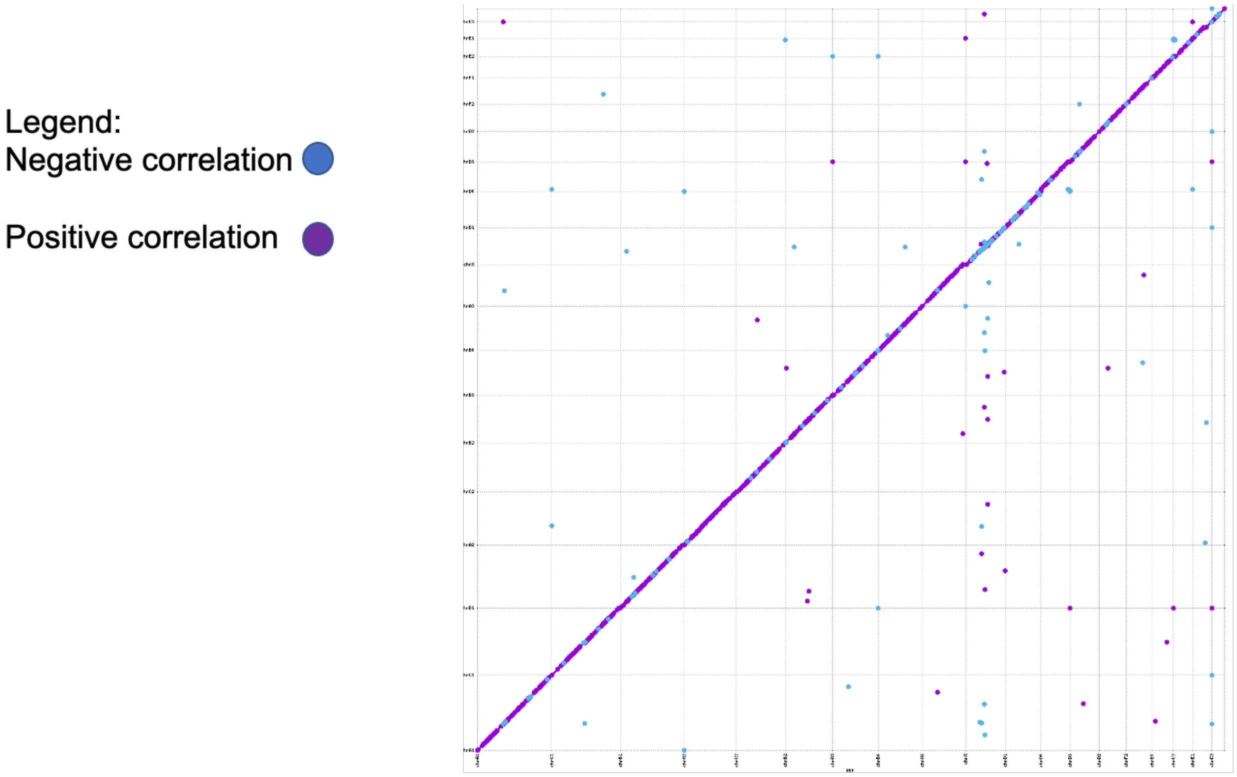


**Supplementary figure 3: Alignment relationship between the Asian leopard cat and fishing cat genome.** Above is the MashMap output comparing the fishing cat chromosome level assembly (y) to the Asian leopard cat assembly (x). Negative alignment correlations are indicated in blue and positive correlations are indicated by purple.

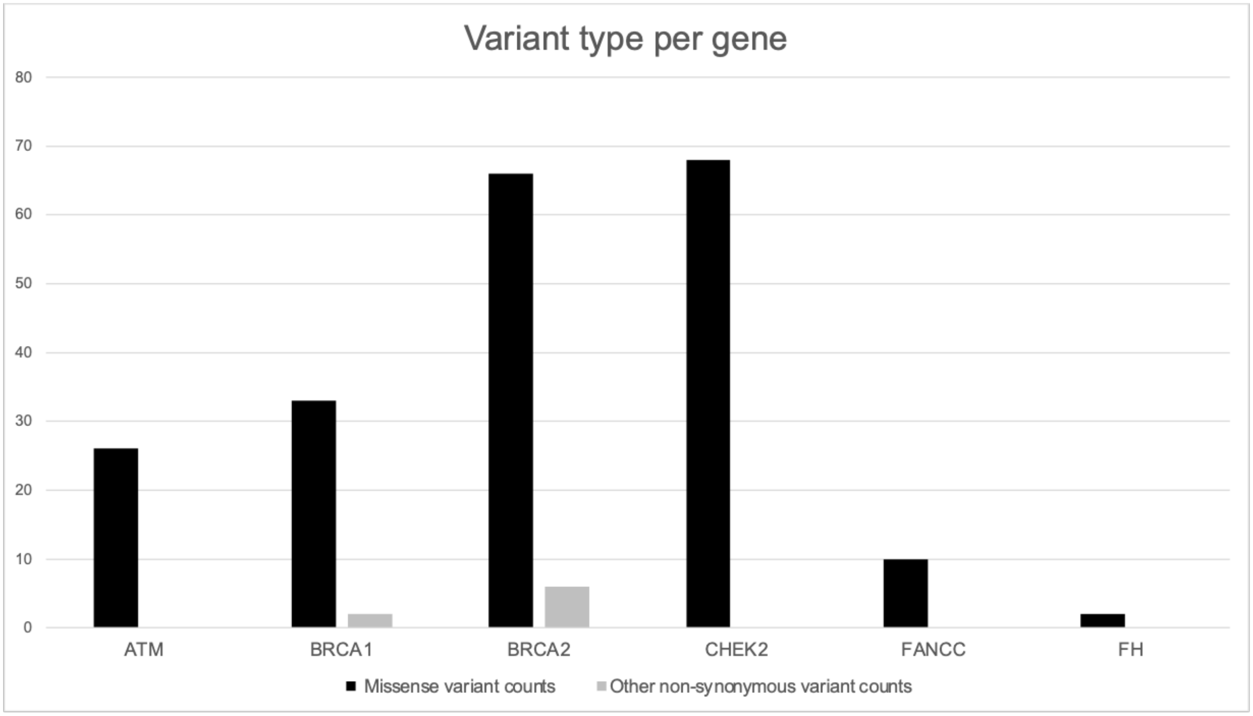


**Supplementary figure 4: Total nonsense variation in fishing cat bladder cancer risk genes.** This bar graph shows the number of missense (black) and other non-synonymous variants located within each gene of interest in the fishing cat cohort. The genes with the most missense variants are *BRCA2* and *CHEK2,* and the only genes with non-synonymous variants are *BRCA1* and *BRCA2*. *BRCA1* has two inframe insertions, and *BRCA2* has 6 inframe deletions.

a.

b.

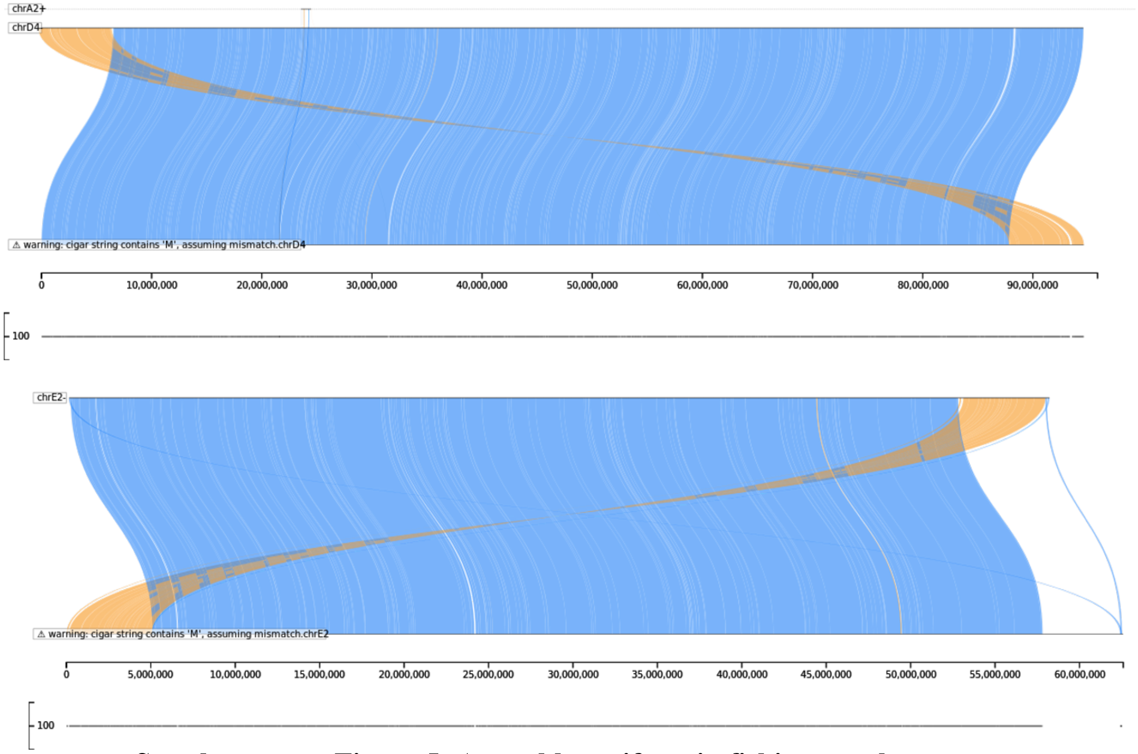


**Supplementary Figure 5: Assembly artifacts in fishing cat chromosomes compared to the reference Asian leopard cat.**

a. Pictured here is the SafFire alignment output between chromosome D4 on the Asian leopard cat (top alignment) and fishing cat (lower alignment).

b. Pictured here is the SafFire alignment output between chromosome E2 on the Asian leopard cat (top alignment) and fishing cat (lower alignment).
